# Data-driven inference of digital twins for high-throughput phenotyping of motile and light-responsive microorganisms

**DOI:** 10.1101/2025.02.21.639423

**Authors:** Andrea Giusti, Davide Salzano, Mario di Bernardo, Thomas E. Gorochowski

## Abstract

Light can affect the movement of microorganisms. These responses can drive collective behaviours like photoaccumulation and photodispersion, which play a key role in broader biological functions like photosynthesis. Our understanding of these emergent phenomena is severely limited by difficulties in obtaining data needed to establish accurate models that can serve as a basis for multi-scale analyses. Here, we address this issue by developing an integrated experimental and computational platform to collect large temporal imaging datasets that allow for the inference of ‘digital twins’ — mathematically precise computational models that accurately mirror the behavior of individual microorganisms — and show that they can replicate the light response of diverse microorganisms *in silico*. We show that a generalised phenomenological model of movement can be effectively parameterised from experimental data to capture key behavioural traits of two commonly studied photo-responsive microorganisms (*Euglena gracilis* and *Volvox aureus*) and demonstrate our model’s ability to accurate reproduce patterns of movement for individuals and populations in response to dynamic and spatially varying light patterns. This work takes steps towards the automated phenotyping of multi-scale behaviours in biology and complements high-throughput genome sequencing efforts by allowing for more comprehensive and quantitative genotype-to-phenotype mappings. It also unlocks new opportunities for the design of spatial control algorithms to guide collective microorganism behaviour.

## INTRODUCTION

Many microorganisms are able to sense light and adapt their movement in response [1–5]. This behaviour is known to be essential for the optimization of photosynthesis [6], minimisation of photodamage [6–8], and is often a key regulator of other cellular processes (e.g., switching of feeding modes or shifts in metabolic states) [9, 10]. While such behaviours are common in nature, how different microorganisms respond to light can vary substantially. Some change their movement speed (photokinesis) [11, 12] or turning rate (photoklinokinesis) [13], while others align to unidirectional light sources (phototaxis) [12, 14] or react to changes in light intensity (photophobic response) [6]. Furthermore, many of these behaviours are observed simultaneously and to differing degrees, and each microorganism response may change over time [9, 15], or be tuned to specific light intensities, wavelengths [16] or even light polarizations [17]. These changes also influence emergent population-level behaviours [18], such as the accumulation of organisms in either illuminated (photoaccumulation) or darker (photodisperison) areas.

While these diverse light-responsive behaviors are widely observed across species, their underlying mechanisms and genetic basis remain poorly understood. This limited mechanistic understanding stems from challenges in collecting the data necessary to dissect responses and create models able to fit experimental observations. Being able to more fully characterize the movement of microorganisms is key to overcoming these difficulties and would help reveal the biological mechanisms behind the regulation of photomovement [2]. Furthermore, the ability to predict how microorganisms respond to light would pave the way for rapidly prototyping light-based controllers to shape emergent collective behaviours for biotechnology applications [19].

A promising avenue for better understanding these complex multi-scale systems is the development of what are termed ‘digital twins’. These are mathematical models that aim to replicate a real-world system *in silico* such that computational simulations can be used to accurately predict responses without the need for time-consuming and laborious experiments. Digital twins for motile microorganisms are primarily constructed in two different ways. First, a bottom-up biochemical and biophysical model can be developed to capture how low-level features like gene expression, biophysics of cilia/flagella, and how shape of the body contribute to changes in movement [20]. Although this approach may allow for a more precise physically-oriented understanding, the resulting digital twin is typically so specific that it cannot be generalised to even closely related species and parameterisation often require biophysical knowledge we do not possess. Furthermore, these models have a high computational complexity that limits scalability and makes it challenging to study emergent population-level behaviours. An alternative approach is to capture the movement of a microorganism by constructing a phenomenological model that is able to describe overall movements in terms of a set of generalised behaviours [21]. Although this approach does not replicate the underlying physiology of the cell, it can efficiently capture the kinematic features of a microorganism’s movement, allowing for the systematic assessment of responses to complex stimuli or simulation of large populations to explore emergent collective behaviours.

Over the past few decades many different phenomenological models have been proposed to describe the locomotion of swimming microorganisms. For example, movement of a wide class of microorganisms, such as *V. aureus, E. gracilis, Chlamydomonas nivalis, Paramecium Bursaria* and *Paramecium Cuadatum* have been described using a Persistent Turning Walker (PTW) model [22–24]. In the simplest version of this model, the longitudinal speed of the microorganism is assumed to be constant, while a stochastic differential equation (SDE) in the form of an Ornstein-Uhlenbeck process (also known as a Vasicek model) [25] describes the evolution of the angular velocity. Extensions of this model have been proposed to capture speed variations in the form of a second Ornstein-Uhlenbeck process, allowing for coupling between the speed and the angular velocity [26], and other extensions have been developed to capture the effects of light [27] on the angular velocity. While the proposed models are able to describe the movement of specific microorganisms, to the best of our knowledge there is no phenomenological model simultaneously accounting for dynamical effects of external inputs (i.e. light) and a non-constant speed. Therefore, none of the existing models allows to reproduce some crucial aspects of photomovement, such as fluctuations in velocity and photokinesis.

Another major challenge when developing digital twins of motile microorganisms is the collection of sufficient data to capture their often noisy dynamics. To characterize the behaviour of a given species, it is typically necessary to conduct large numbers of experiments in which the microorganisms are exposed to a variety of lighting conditions. Although some platforms have been proposed to image and stimulate light sensitive microorganisms[28–30], these lack the ability to perform semi-automated characterization of microorganisms in both space and time.

In this work, we aim to address these challenges by developing an integrated low-cost platform to collect and process imaging data sets that can be used to perform data-driven inference of digital twins for motile microorganisms. We extend the hardware and software of an existing imaging and light projection system called the Dynamic Optical MicroEnvironment (DOME) [30, 31] and create a generalised mathematical model able to capture the light response of diverse types of microorganism. We use this platform to rapidly infer digital twins for *E. gracilis* and *V. aureus*, and show the ability for our models to accurately replicate single cell behaviours and emergent population level responses seen in experiments not used for model fitting. Overall, this work advances the high-throughput quantitative phenotyping of motile and light-responsive microorganisms by providing an integrated platform that bridges individual and population-level behaviors. The combination of automated experimental characterization with accurate digital twins enables systematic investigation of complex multi-scale biological phenomena and opens new possibilities for designing light-based control strategies in biotechnology applications.

## RESULTS

### Platform for inferring digital twins of motile microorganisms

We began by developing an integrated platform for the data-driven generation of digital twins (**Figure 1**). The physical platform was based on the DOME [30, 31]; a low cost, modular, and open-source experimental system capable of imaging living organisms in real-time and simultaneously projecting dynamic light patterns across a sample at resolutions that enabled illumination of individuals in a population. We extended the DOME’s software architecture by adding new modules to control the imaging and light projection systems, as well as developing new analysis modules to process the imaging data produced (**Methods**). These extensions allowed us to run experiments with light inputs that varied both in space and time, and to automate the data processing.

**Figure 1:**
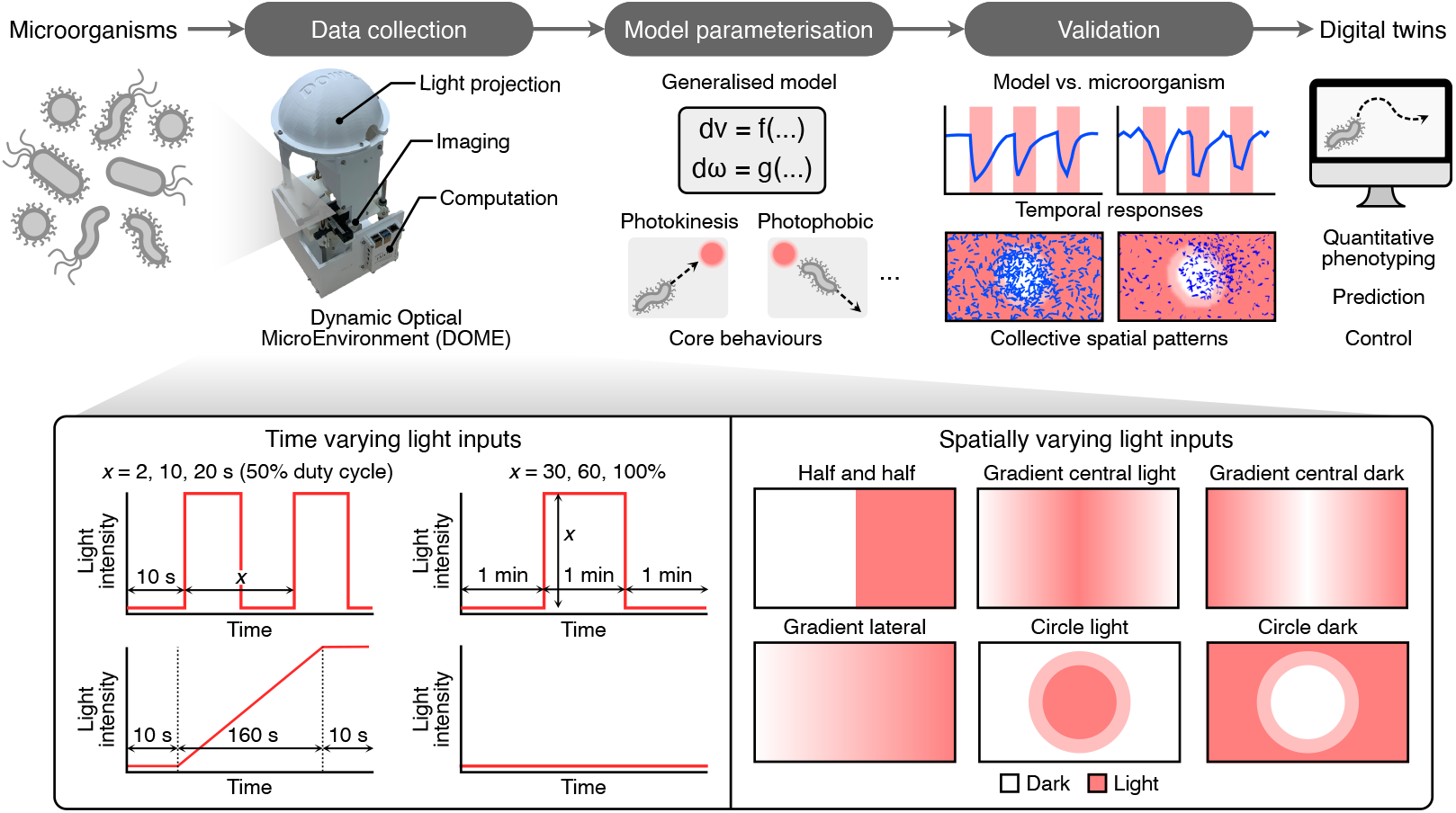
Overview of the high-throughput phenotyping system for motile and light responsive microorganisms. The platform consists of three major steps. First, data collection is performed using the Dynamical Optical MicroEnvironment (DOME) system that enables the imaging and illumination of microorganisms over time. During this stage, we expose microorganisms to a range of temporally and spatially varying light patterns (bottom box) and monitor their response. The platform includes data processing modules to generate movement speeds and directions for all microorganisms present over time. The second stage is model parameterisation that uses a generalised model of microorganism movement and response to light in conjunction with the characterisation data to fit refined models that fit the population-level variation we observed during characterisation. Finally, the third stage validates the accuracy of the model, using characterisation data that was not used for model fitting. Overall, the platform enables us to take a microorganism of interest and then rapidly generate digital twins that are able to replicate core light-responses and movement.

The generation of a digital twin involves three major steps. In the first step, we collect movement data by exposing the microorganism to a standardized set of spatio-temporal light patterns designed to probe their full range of responses (**Methods**). This standardization ensures we capture sufficient information to robustly fit our mathematical model parameters. A critical challenge during this step is accurately tracking individual microorganisms. This involves the removal of static background features, identification of microorganism positions using a threshold-based method, and the application of a tracking algorithm to detect and label microorganisms across frames (**Methods** and **Supplementary Figure 1**). To improve the quality of the recovered trajectories, a multi-step refinement algorithm is also used to attenuate measurement noise. This processing pipeline enables accurate frame-by-frame calculation of each microorganism’s speed and angular velocity.

Once the trajectories have been recovered for a broad range of light inputs, the second step consists in the creation of digital twins, by fitting our generalised model to experimental data. Statistical analysis of the experimental trajectories is used to tailor the model (i.e., select the relevant core behaviours) of the observed microorganism and then, for each microorganism in the population, a parameter set is estimated using a multi-step identification procedure (**Methods**). Crucially, this approach captures the variability of parameters across the population, reproducing the specific behaviours of specimens.

The third and final stage consists in the validation of the model. This is performed by using our digital twin as a basis for agent-based simulations. By comparing our *in silico* simulations to real-world experiments for different light inputs, varying in space and time (which have not been used for fitting the model), we can quantitatively assess the ability for the digital twins to reproduce behaviours observed at the level of single microorganisms and the level of the collective.

### A generalised model of motile and light responsive microorganisms

Central to our approach is the ability to capture the response of a microorganism to light inputs using a mathematical model. Although microorganisms can respond to light in many different ways, these phenotypes can often be decomposed into a small set of core behaviours (e.g., photokinesis, photoklinokinesis, and step-up and step-down photophobic responses) that can be modelled separately and then combined. Using this approach, we developed a generalised phenomenological model (an extension of the PTW model [22]) that is able to capture key kinematic features exhibited by light sensitive microorganisms. Our model consists of two stochastic differential equations that describe the evolution of the longitudinal speed *v* and the angular velocity *ω* in response to a light stimulus *u*. Considering a population of microorganisms, the dynamics of the *i*-th individual are described using:

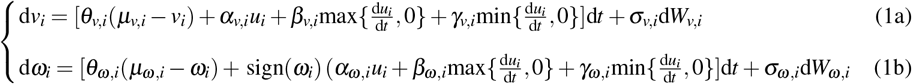

Here, *µ*_*v,i*_ and *µ*_*ω,i*_ are the average longitudinal and angular velocities, *θ*_*v,i*_ and *θ*_*ω,i*_ are the convergence rates of *v* and *ω, σ*_*v,i*_ and *σ*_*ω,i*_ represent the volatilities, and *W*_*v,i*_ and *W*_*ω,i*_ are independent standard Wiener processes. The remaining parameters describe how the microorganism responds to light. Specifically, *α*_*v,i*_ and *α*_*ω,i*_ modulate the intensity of photokinesis and photoklinokinesis, respectively, while *β*_*v,i*_ and *β*_*ω,i*_, and *γ*_*v,i*_ and *γ*_*ω,i*_ capture step-up and step-down photophobic responses. It should be noted that 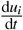 represents the weak derivative of *u*, allowing for discontinuities in the input signal and modelling the effect that changes in the light input have on the microorganism dynamics.

In the absence of a light stimulus, each microorganism on average converges with time constants *θ*_*v,i*_, *θ*_*ω,i*_ to a constant speed *µ*_*v,i*_ and angular velocity *µ*_*ω,i*_. This behaviour can be modified by illuminating the microorganism. In the case where *α*_*v,i*_*≠* 0 (i.e., photokinesis), the average steady-state of *v*_*i*_ changes linearly with the light intensity *u*_*i*_. Similarly, if the microorganism exhibits photoklinokinesis (*α*_*ω,i*_*≠* 0), then the average steady-state of |*ω*_*i*_| varies linearly with the amplitude of the light source. Furthermore, if *β*_*v,i*_*≠* 0 and *β*_*ω,i*_*≠* 0 then a step-up photophobic response is observed, meaning that *v*_*i*_ and |*ω*_*i*_| changes in response to an increase in light intensity. Finally, if *γ*_*v,i*_*≠* 0 and *γ*_*ω,i*_*≠* 0 then a step-down photophobic response is displayed, resulting in a change in *v*_*i*_ and |*ω*_*i*_| when there is a decrease in light intensity. The function sign(*ω*_*i*_) multiplying the input terms in Equation (1b) was added so that the sign of 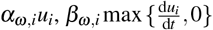 and *γ*_*ω,i*_ min 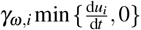 defines the sign of the variation of |*ω*_*i*_| (see **Methods** for more details on the evolution of |*ω*_*i*_|). Specifically, this choice ensures that positive values of these terms correspond to an increase in the magnitude of the angular velocity, irrespective of the current turning direction of the microorganism. Conversely, a negative value corresponds to a decrease in |*ω*_*i*_|.

### Data-driven inference of a digital twin for *Euglena gracilis*

To assess the ability for our platform to efficiently infer digital twins of light-responsive microorganisms, we began by focusing on *E. gracilis*. This is a widely studied aquatic microorganism that is commonly used to study photosynthesis and photoreception, as well as recently being engineered to act as a basis for non-animal based foods [32]. We performed three independent experiments for each input profile, to ensure that we captured both intra- and inter-experiment variability. Data from these experiments were then aggregated to increase the data available for the inference of model parameters (**Methods**).

We started by investigating the movement of *E. gracilis* in a dark environment. We observed an average speed of 60 µm/s, which remained approximately constant during the duration of the experiment (**Figure 2a**). We also analysed the distribution of angular velocities (over time) across the population (**Figure 2b**). Contrarily to what has been previously reported [27, 33], we did not observe a preferential direction of rotation, with a median value that did not statistically deviate from 0 (sign-test, *p* = 0.27; **Figure 2c**).

**Figure 2:**
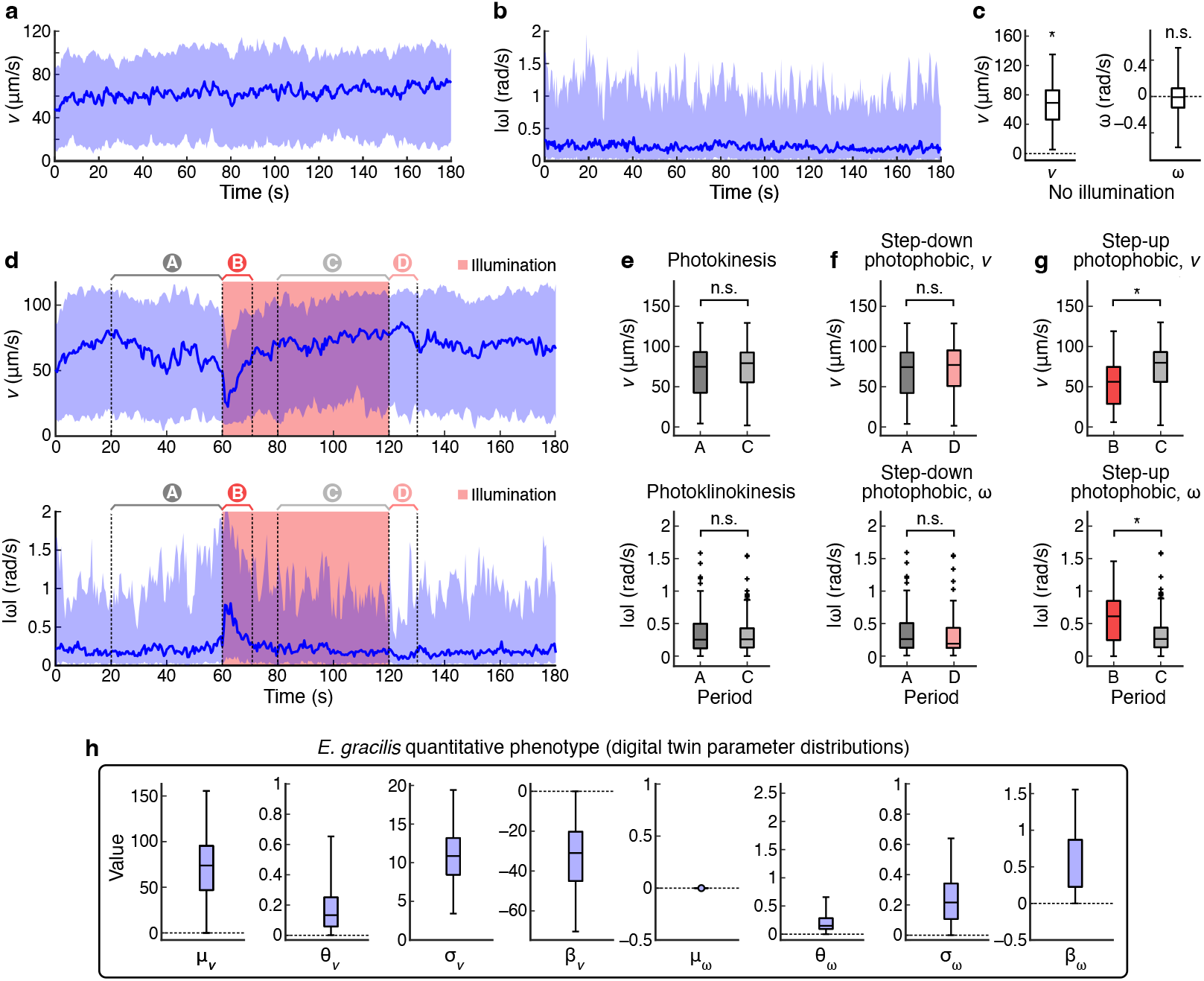
Data-driven phenotyping of *E. gracilis*. (**a**) Time evolution of the longitudinal velocity, *v*, in a dark environment. (**b**) Time evolution of the absolute angular velocity, |*ω*|, in a dark environment. Solid lines represent the median over the population while the shaded areas denotes the 10th-90th percentile range. (**c**) Distributions of the average longitudinal velocity and average angular velocity for the experiment shown in panel **a**. A sign-test was used to test the hypothesis that the distribution has null median. (**d**) Time evolution of *v* (top) and |*ω*| (bottom) when the microorganisms were stimulated using a square wave light input with semi-period of 60 s. The solid lines represent the median over the population, and the shaded areas highlight the 10th-90th percentile range. The red segments highlights the interval when light was present. The highlighted regions A, B, C and D capture regions before and after illumination to enable characterisation of core behavioural responses. Comparison of the distributions of average *v* (top) and |*ω*| (bottom) in selected time windows A–D to assess the presence of (**e**) photokinesis/photoklinokinesis, (**f**) step-down photophobic responses, and (**g**) step-up photophobic responses. The difference between the distributions in each panel was quantified using a Kolmogorov-Smirnov test (n.s. denotes not significant; * denotes statistical significance with *p <* 0.001). (**h**) Distributions of model parameters across the *E. gracilis* population resulting from the identification procedure.

Next, we characterised the response of *E. gracilis* to a variety of light inputs. We specifically ran sets of experiments with spatially uniform illumination across the sample, but where the intensity of illumination varied over time as a square wave with a duty cycle of 50% and a semi-period ranging from 1–60 s (**Methods**). We observed a strong step-up photo-phobic response to this input, followed by a slow adaptation to the increased light intensity with an average adaptation time of ∼10 s. Only in the case of a fast switching square wave (1 s semi-period) we did not observe any significant response (**Supplementary Figure 2**), suggesting an adaptation to (or filtering of) fast switching stimuli.

Given this observed behavior, we selected a square wave input with a 60 s semi-period to quantitatively characterize the types of light response exhibited. This external input guarantees that steady state is reached both in presence and in absence of light. We tested for the presence of photokinesis and photoklinokinesis by comparing the distribution of the average speed and absolute angular velocity of the microorganisms 40 s before the light switches off and 40 s before the light switches on (**Figure 2d**). In both cases, distributions were not statistically different (Kolmogorov-Smirnov test, *p* = 0.775 for *v*, and *p* = 0.365 for |*ω*|), suggesting that neither photokinesis nor photoklinokinesis occurs (**Figure 2e**). This is further supported by the lack of response observed when changing the light input slowly, for example, by linearly increasing the light intensity from its minimum to maximum value (**Supplementary Figure 3**). To test for the presence of a step-down photophobic response, we compared the distributions of the average *v* and |*ω*| across the population 40 s before the light switches on and 12 s after the light switches off. These distributions were not statistically different both for *v* (*p* = 0.46) and |*ω*| (*p* = 0.12), suggesting the absence of this behaviour (**Figure 2f**). Finally, we tested for presence of a step-up photophobic response by comparing the distributions of the average speed and angular velocity of the agents 40 s before the light switches off and 12 s after the light switches on. In this case, distributions of *v* and |*ω*| were both statistically different in the two lighting conditions (*p* = 2.16 *×* 10^−6^ and *p* = 1.07 *×* 10^−8^, respectively), confirming the presence of a strong step-up photophobic response (**Figure 2g**).

To complete the characterisation, we validated the modelling assumption that the increase/decrease in *v* and |*ω*| scales linearly with the derivative of the light intensity by performing experiments with square waves with fixed 60 s semi-period, but different intensities of illumination. By fitting a linear regressor to the average fold-change of *v* and |*ω*| for the first 12 s after the light switches on, with respect to the average 40 s before the light switches on, we obtained a *R*^2^ = 0.62 for *v* and *R*^2^ = 0.98 for |*ω*|, confirming the validity of our modelling assumption (**Supplementary Figure 4**).

From these analyses it was possible to define a simplified model for *E. gracilis*, by omitting terms describing behaviours not seen in our experimental data (i.e., *µ*_*ω,i*_ = *α*_*v,i*_ = *α*_*ω,i*_ = *γ*_*v,i*_ = *γ*_*ω,i*_ = 0). After these simplifications, the movement and response of an *E. gracilis* microorganism could be described using

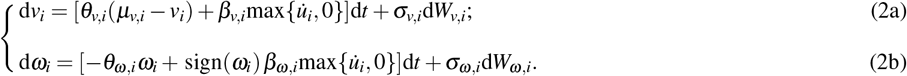

In addition, as shown in **Figure 2f**, positive changes in light intensity cause an increment of |*ω*| and a decrease of *v*. Therefore, we imposed *β*_*ω,i*_ *>* 0 and *β*_*v,i*_ *<* 0. These constraints, together with the assumption of stable dynamics (i.e., *θ*_*v,i*_, *θ*_*ω, i*_ *>* 0), were used to constrain the parametrization.

Having inferred the structure of our digital twin, the final step was to identify appropriate parameters to replicate experimentally measured behaviours. The parametric identification was performed using data from the experiment with square wave input of 10 s semi-period. This input was chosen to induce photophobic responses repeatedly, while giving the organisms enough time to adapt after each change of light intensity. For each specimen in the population, parametrization was performed independently for *v* and *ω*, using a three steps procedure. Note that this is possible since equations (2a) and (2b) are decoupled. First we identified the set of parameters fitting the average behaviour of the population at each time point, combining ordinary least squares (OLS) with the MATLAB GrayBox identification toolbox. This parameter set was used as an initial guess to estimate, for each microorganism within the population, the value of the parameters fitting its longitudinal and angular velocity profiles. This allowed the digital twin to replicate the observed kinematic features. Finally, outlier digital twins were identified and removed (**Methods**). The resulting distributions of the model parameters are shown in **Figure 2h**. By generating a different set of parameters for each specimen, we were able to capture the intrinsic heterogeneity of the population while preserving correlations between the parameters.

### Validation of the *Euglena gracilis* digital twin

The analysis of the experimental trajectories together with the identification procedure allowed for the generation of digital twins that quantitatively replicated the movement of *E. gracilis*. To assess the ability of our digital twins to reproduce the movement of real microorganisms, we mimicked *in silico* each experimental conditions tested above using the agent-based simulation tool SwarmSim (**Methods**). Specifically, we replicated each experimental condition by using the same light stimuli and simulating the movement of the digital agents identified from experimental data.

For each experiment, we assessed the ability of the digital twins to reproduce the experimentally observed behaviour. We found a good qualitative agreement between the experimental data and their digital counterpart in most scenarios (**Figure 3a–h**). The digital twins reproduce both the steady state behaviour of the microorganisms and their step-up photophobic response followed by adaptation to the new condition. To measure the accuracy of our digital twins, we evaluated the discrepancy between the median of simulated and experimental data using the average weighted mean absolute percentage error (wMAPE). This metric assumes similar values across most scenarios tested, highlighting the ability of our digital twins to reproduce the observed photo-movement in experimental conditions that were not observed during parameter fitting (**Figure 3i**). The only condition we were not able to replicate was the response to a square wave with a 1 s semi-period. In this scenario, our model significantly overestimates the response of the agents to the light stimuli (**Figure 3a**). This may be caused by the inability of our simple first-order dynamics to efficiently filter out inputs at such high frequencies.

**Figure 3:**
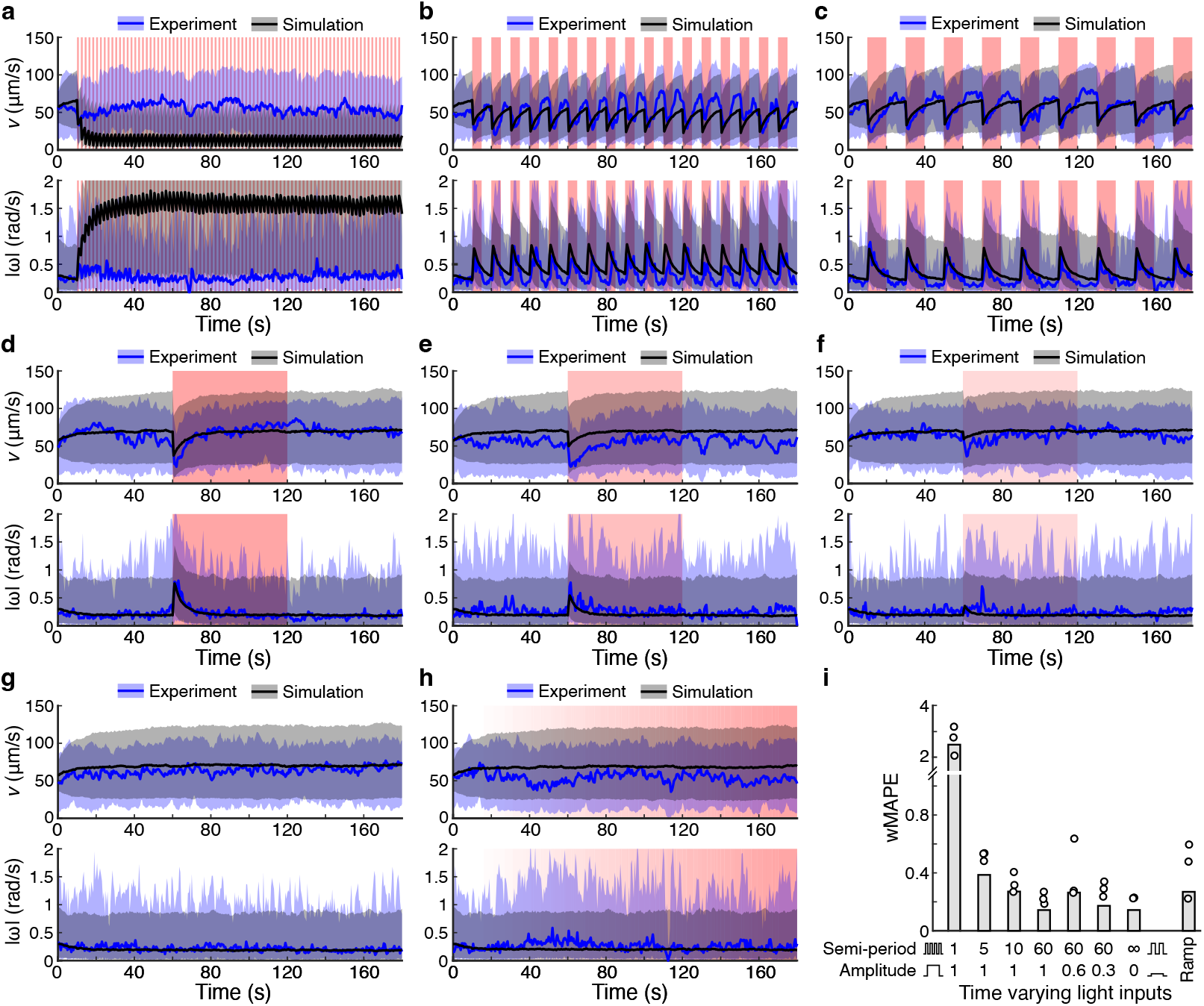
Validation of generalised *E. gracilis* model using time varying light inputs. Comparisons of the time evolution of longitudinal velocity *v*, and absolute angular velocity |*ω*| (bottom), between experiments (combined biological replicates, blue) and model simulations (black) with different temporal light patterns. Solid lines represent the median over the population while the shaded areas highlight the 10th-90th percentile range. Red shaded areas highlight the intervals when light inputs were present. (**a**) Square wave light input, 1 s semi-period, 100% intensity. (**b**) Square wave light input, 5 s semi-period, 100% intensity. (**c**) Square wave light input, 10 s semi-period, 100% intensity (used for parameters identification). (**d**) Square wave light input, 60 s semi-period, 100% intensity (used for model selection). (**e**) Square wave light input, 60 s semi-period, 60% intensity. (**f**) Square wave light input, 60 s semi-period, 30% intensity. (**g**) No illumination. (**h**) Linearly increasing light intensity input from 0% to 100%. (**i**) Average weighted mean absolute percentage error (wMAPE) for different experimental conditions. The wMAPE metric was computed by comparing the median of simulation outputs to the median of the experiments where data for all biological replicates was combined. Dots represent the wMAPE when comparing simulations to each biological replicate separately.

Having shown the ability of our digital twins to reproduce the dynamic evolution of *v* and *ω* in time, we next tested whether they are able to show the same photodispersion exhibited by *E. gracilis* when exposed to spatially varying light inputs. We performed experiments where a static spatial input was shed on to a *E. gracilis* population for 180 s. We specifically projected the letters BCL (representing the initials of the experimental lab) as dark regions oven an illuminated background, and observed accumulation of microorganisms in the darker areas (**Supplementary Movies 1 and 2**). We repeated the same experiment *in silico* using SwarmSim and observed that the digital twins accumulating in the same areas (**Figure 4a**).

**Figure 4:**
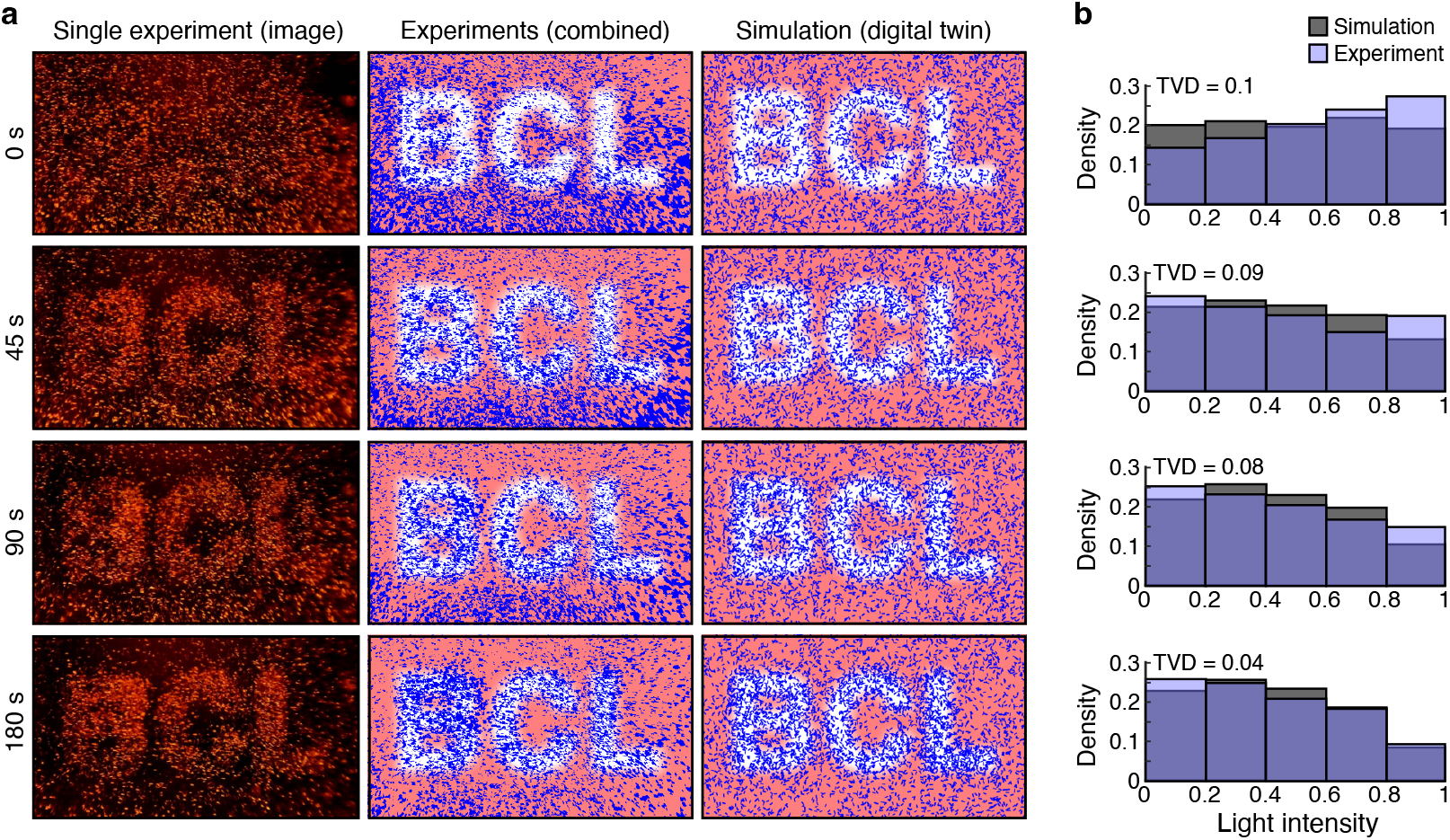
Response of *E. gracilis* to a complex spatial light pattern. A static light pattern showing the letters ‘BCL’ (in reference to the name of our experimental lab) was provided as input to a population of *E. gracilis* and induces photodispersion. (**a**) Snapshots of the spatial organisation of the population at different time points: (left column) raw microcopy images from the DOME, (middle column) combined data from experiments, (right column) simulation data. The red shaded pattern shows the light input projected onto a dark/unilluminated background shown in white. (**b**) Density distributions of the microorganisms as a function of the normalized light intensity (simulations: dark grey; experiments: light purple). The total variation distance (TVD) between the simulations and experiments is shown for each plot.

To better understand this response, we quantified the degree of photodispersion by estimating the density distribution of the microorganisms as a function of the normalized light intensity (**Methods**). Using the histograms from experiments and *in silico* simulations (overlaid in **Figure 4b**), it is possible to see that in both scenarios the microorganisms progressively accumulate in the areas of lower light intensity. To measure the difference between the density distributions of experiments and simulations, we calculated the Total Variation Distance (TVD) (**Methods**). Throughout the all experiment, the TVD never exceeded 0.1, confirming the ability of the digital twins to reproduce the spatial patterns that emerged.

Similar comparisons were also performed for a broader range of spatially varying inputs (**Figure 5a–f**). For all six scenarios we considered, simulations of the digital twin qualitatively reproduced the experimental data, exhibiting different degrees of photodispersion depending on the input light pattern. In addition, although experimental data showed a more pronounced photodispersion when exposed to some light patterns, the TVD between simulation and experimental density distributions never exceeded 0.5 and was typically below 0.3 (**Figure 5g**). This provides further support that our digital twins are able to replicate diverse photo-induced behaviours of *E. gracilis*.

**Figure 5:**
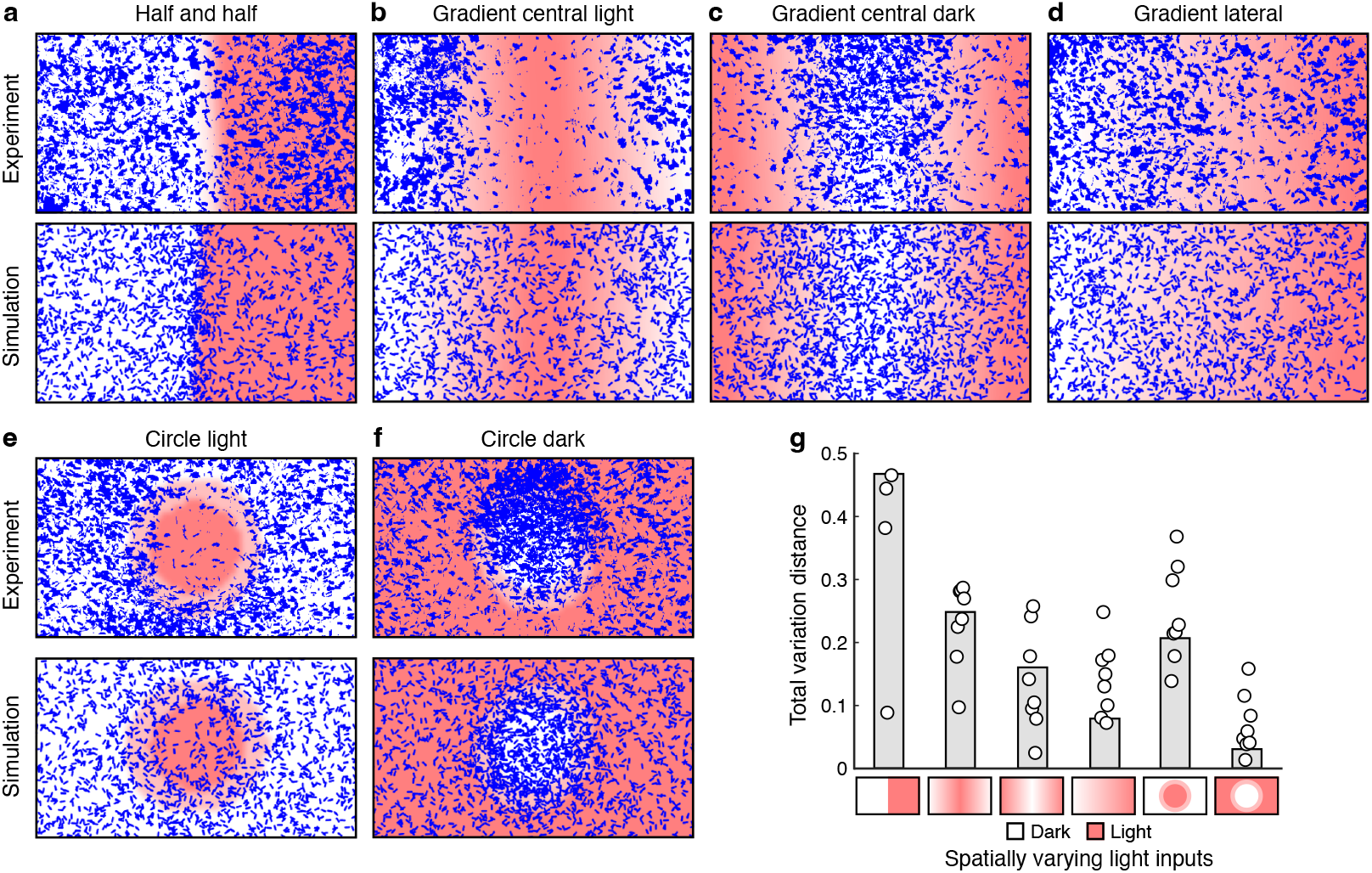
Validation of generalised *E. gracilis* model using spatially varying light inputs. Snapshots from experiments and simulations after 180 s of exposure to a spatially patterned light inputs. Red and white regions denote illuminated and dark/unilluminated areas, respectively. (**a**) Half and half light pattern. (**b**) Gradient from lit centre. (**c**) Gradient from unlit centre. (**d**) Lateral gradient dark to light (left to right). (**e**) Central lit circle and unlit background. (**f**) Central unlit circle and lit background. (**g**) Total variation distance (TVD) between experimental and simulated density distributions of the microorganisms as a function of the normalized light intensity. The TVD metric was computed by comparing simulation outputs to the combination of all biological replicates. Dots represent the TVD when comparing simulations to each biological replicate separately.

### Generality of the phenotyping platform and model

To ensure that our approach could be applied to other types of microorganism, we carried out further experiments to infer digital twins for *Volvox aureus. V. aureus* is a micro-algae that forms large spherical multi-cellular colonies up to 1 mm in diameter, which are both motile and light responsive. It is also known that the type of motility and response of *V. aureus* is very different to *E. gracilis* due to mechanistic differences in how motion is achieved [34], further assessing the flexibility of our generalised model.

Due to the size of the *V. aureus* colonies, we needed to adapt the DOME to use a lower magnification objective for imaging (**Methods**). Similar experiments as for *E. gracilis* were then performed to characterize the response of *V. aureus* to various types of illumination. In a dark environment, *V. aureus* moved at an average longitudinal speed of 40 µm/s (**Figure 6a**). Unlike *E. gracilis, V. aureus* showed a significant preference for turning in an anticlockwise direction (**Figure 6b**), with a median average angular velocity of 0.134 rad/s (sign-test, *p* = 2.1 *×* 10^−10^; **Figure 6c**). Temporally varying light inputs showed that *V. aureus* required ∼20 s to adapt to a new light intensity (**Figure 6d**). Furthermore, a strong step-up photophobic response for *v* appeared as the only light-dependent behaviour seen (**Figure 6e–g**). While no statistically significant response for |*ω*| was observed.

**Figure 6:**
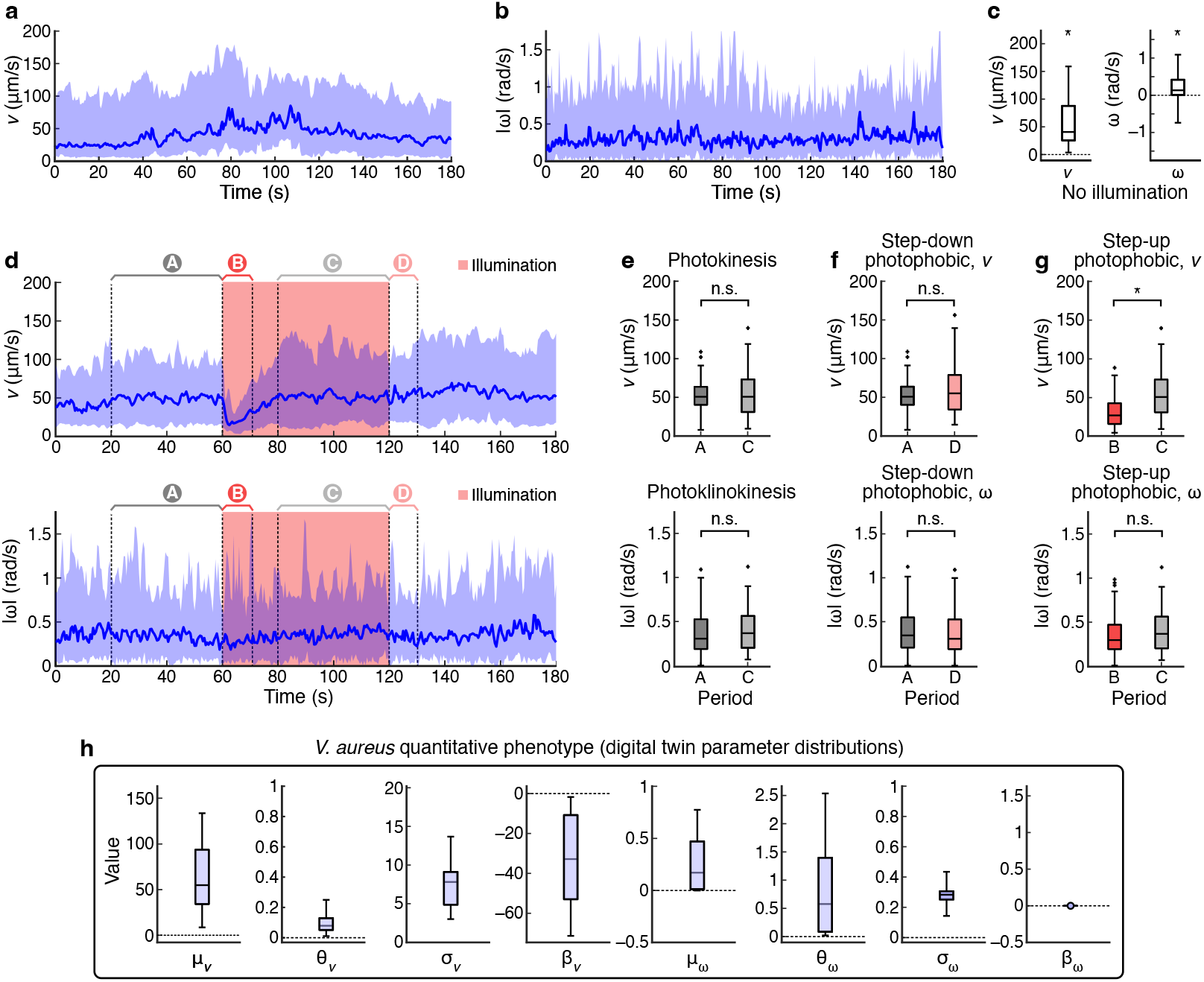
Data-driven phenotyping of *V. aureus*. (**a**) Time evolution of the longitudinal velocity, *v*, in a dark unilluminated environment. (**b**) Time evolution of the absolute angular velocity, |*ω*|, in a dark unilluminated environment. Solid lines represent the median over the population while the shaded areas denotes the 10th-90th percentile range. (**c**) Distributions of the average longitudinal velocity and average angular velocity for the experiment shown in panel **a**. A sign-test was used to test the hypothesis that the distributions have null median. (**d**) Time evolution of *v* (top) and |*ω*| (bottom) when the microorganisms were stimulated using a square wave light input with semi-period of 60 s. The solid lines represent the median over the population, and the shaded areas highlight the 10th-90th percentile range. The red segments highlight the interval when light was present. The highlighted regions A, B, C and D capture regions before and after illumination to enable characterisation of core behavioural responses. Comparison of the distributions of average *v* (top) and |*ω*| (bottom) in selected time windows A–D to assess the presence of (**e**) photokinesis/photoklinokinesis, (**f**) step-down photophobic responses, and (**g**) step-up photophobic responses. The difference between the distributions in each panel was quantified using a Kolmogorov-Smirnov test (n.s. denotes not significant; * denotes statistical significance with *p <* 0.001). (**h**) Distributions of generalised model parameters across the *V. aureus* population that are used to generate digital twins.

Based on the strong step-up photophobic response, it was possible to generate a simplified model for *V. aureus* movement and its response to light (i.e., *α*_*v,i*_ = *α*_*ω,i*_ = *β*_*ω,i*_ = *γ*_*v,i*_ = *γ*_*ω,i*_ = 0). Specifically,

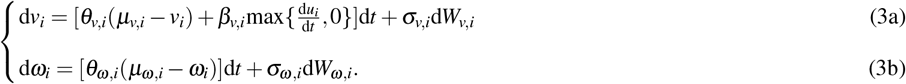

This model was then parameterised using data from square wave input experiments with a semi-period of 60 s. This input was chosen to induce a photophobic response, while giving the microorganisms sufficient time to adapt to the new light intensity. For each individual in the experimental population, parameters were inferred using the same approach as for *E. gracilis*, resulting in the distributions shown in **Figure 6h**.

As before, we validated the accuracy of our digital twins by comparing experiments with simulations for a range of uniform and time varying light inputs (**Supplementary Figure 5**). For both of these, our digital twins were able to accurately reproduce the experimental data, obtaining similar wMAPE in virtually all conditions (**Figure 7a**). The only exception was for a square wave with a 1 s semi-period, where similarly to *E. gracilis*, our digital twins overestimated the variation of the longitudinal speed.

**Figure 7:**
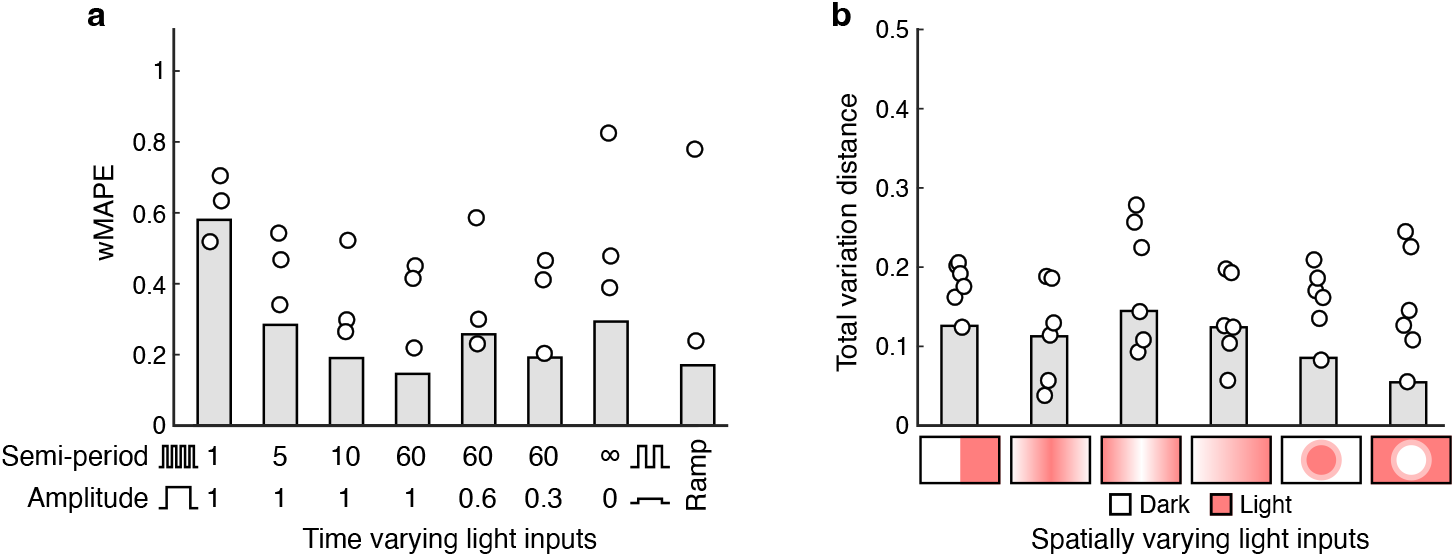
Validation of generalised *V. aureus* model using temporally and spatially varying light inputs. (**a**) Average weighted mean absolute percentage error (wMAPE) for different experimental conditions with temporally varying light inputs. The wMAPE metric was computed by comparing median of the simulation outputs with the median of the experiments where data for all three biological replicates was combined. Dots represent the wMAPE values when comparing simulations to each biological replicate separately. (**b**) Total variation distance (TVD) in density distributions of the microorganisms between experiments and simulations as a function of the normalized light intensity. The TVD metric was computed by comparing simulation outputs to the combined biological replicates. Dots represent the TVD when comparing simulations to each biological replicate separately.

Finally, we tested spatially varying light inputs. Both experiments and simulations showed a weak photoaccumulation (**Supplementary Figure 6**), which differed substantially from the behaviour of *E. gracilis*. We compared the steady state distributions for each of these input types and found a good qualitative agreement (**Supplementary Figure 7**). Furthermore, as for *E. gracilis*, we measured the discrepancy in the distributions using the TVD metric. In all light conditions, the TVD never exceeded 0.3 (**Figure 7b**), confirming that the inferred digital twins were able to accurately reproduce the experimentally observed spatial patterns.

## DISCUSSION

In this work, we developed an integrated experimental and computational platform to enable the rapid inference of digital twins for a wide class of light-responsive microorganisms. We demonstrated how a single characterisation experiment can be used to uncover key features of photomovement associated with diverse types of microorganism, and how this information can be used to fit parameters of a generalised model capturing quantitative phenotypic information. Furthermore, we show how this model can then be used to accurately replicate complex spatial-temporal behaviours observed in experiments not used for parameterisation of the model.

Our ability to effectively phenotype light-responsive microorganisms was made possible by adapting the DOME platform, which enabled precise dynamic light patterns that would have been difficult to achieve using standard imaging equipment, as well as the development of a generalised model able to more fully capture the range of light responses possible. To the best of our knowledge, our model is the first that is able to describe changes in both longitudinal and angular velocity, as well as capture responses in these variables due to light. Despite its simplicity, our results show that this model is able to reproduce a wide variety of behaviours observed in light sensitive microorganisms. However, since it is designed to describe microorganisms moving on a plane with light shed from a unidirectional source (above the sample stage), it cannot capture effects that depend on the direction of the incoming light (e.g., phototaxis). Furthermore, the model neglects possible interactions between microorganisms, such as collisions or chemical communication, and as shown in our results, struggles to capture high-frequency inputs. Extensions to capture these aspects would be valuable additions to this work.

Our digital twins pave the way to the development of light-based controllers to shape emergent collective behaviours of microorganisms, with relevant applications in biotechnology and synthetic biology [29, 35]. The use of external stimuli to control emergent macroscopic features of cellular populations has been demonstrated with bacteria [36], but extending this approach to more complex microorganisms has seen less attention in part due to limited platforms to carry out such experiments. Moreover, the ability to rapidly prototype new control architectures using our digital twins as a foundation, could also offer new insights into effective controller design for biological systems where noise and heterogeneity are unavoidable. If successful, such controllers could have wide ranging impact in many different areas spanning wound healing [37] to diagnostics and sensing [31, 38]. Examples of potential applications include (i) guiding microalgal populations for optimized biofuel production by controlling their spatial distribution in photobioreactors; (ii) enhancing wastewater treatment by manipulating the movement of photosynthetic microorganisms to increase metabolic efficiency; (iii) improving microscale biosensing by precisely controlling the spatial organization of light-responsive microorganisms; (iv) designing new microfluidic sorting techniques based on phototactic responses Furthermore, the platform could accelerate strain development by enabling rapid phenotypic characterization of engineered variants.

With the phenotyping of microorganisms often acting as a bottleneck in our ability to understand the diversity of microbial behaviours and their possible role in broader ecosystem functions, this work provides a step towards more automated, rapid and quantitative approaches, demonstrating the value of integrating custom designed hardware and generalised model for the biological sciences.

Future development of this platform will focus on several key areas. Integration with automated microscopy systems could enable truly high-throughput phenotyping of large strain libraries. The mathematical model could be extended to incorporate multi-wavelength responses and organism-organism interactions. Addition of real-time tracking and control capabilities would enable closed-loop optimization of light patterns for desired collective behaviors. The platform could also be adapted for other stimuli beyond light, such as chemical gradients or temperature variations.

## METHODS

### Microorganisms and culturing

*V. aureus* and *E. gracilis* strains were obtained from BladesBio UK. *V. aureus*s was cultured in commercial Alga Grow medium (BladesBio UK) following manufacturer protocols. *E. gracilis* was maintained in a defined medium prepared by boiling 40 wheat grains and 35 rice grains in 1 L of deionized water, then filter-sterilizing the resulting solution. Both organisms were cultured at ambient temperature (22 ± 2°C) under full-spectrum artificial illumination with a 12 hour light/dark cycle.

### Experimental setup

All experiments were carried out using the DOME [30] with an updated high-definition (HD) camera [39]. To interact with the DOME, we used a USB keyboard, mouse, and HDMI monitor connected to the Raspberry Pi 4. We used the DOME in three different configurations: (i) with a 10X objective lens (90X total magnification) for experiments with *E. gracilis*; (ii) with a 4X objective lens (36X total magnification) for the BCL experiment with *E. gracilis*; and (iii) without objective lens (9X total magnification), for experiments with *V. aureus*. The liquid culture was directly loaded onto microscopy slides, apart from experiments with *V. aureus* where the slides were fitted with a 3D printed plastic frame [30] to allow for the use of larger liquid samples (up to 300 µL). For standard experiments with *E. gracilis*, 15–20 µL of cultures containing 50–100 cells/µL were used. For the BCL experiment with *E. gracilis*, 20 µL of cultures containing 65– 75 cells/µL were used. For experiments with *V. aureus*, 120–150 µL of cultures containing 0.2–0.8 cells/µL were used.

### Experimental protocols

We performed a set of open loop experiments using either time varying, spatially uniform or spatially defined static light patterns. Each experiment lasted for 180 s and images were taken every 0.5 s. The sampling time was chosen due to constraints on the imaging elements of the DOME, while the duration of the experiment was selected to allow for the observation of slow dynamics, such as adaptation, while containing the cost in terms of time and memory requirements for the DOME.

For each experiment, a small amount of continuous culture was placed on a microscopy slide. The slide was then placed on the sample stage of the DOME and the device covered with a dark hood to filter out any external light. During the entire duration of the experiment the projector was set to emit a constant and low intensity red light (640 nm at 5% of the projector’s maximum brightness). This provided illumination for dark-field imaging. For stimulating the microorganisms, blue light (460 nm) was used. A band pass filter that only allowed for red light frequencies from our projector was used to prevent blue light from reaching the camera sensor and improving the quality of image acquisition.

The following 8 experiments with time-varying inputs were performed: no input illumination; square waves inputs with 100% intensity and 1 s, 5 s, and 10 s semi-periods; square waves inputs with 30%, 60%, and 100% intensity and 60 s semi-period; and linearly increasing intensity from 0% at *t* ≤ 10 s, to 100% intensity for *t* ≥ 170 s (**Figure 1**).

The following 6 experiments with spatially defined inputs were performed: unlit left half and 100% lit right half, linear gradient from lit centre to dark edges, linear gradient from dark centre to lit edges, linear gradient from dark left edge to lit right edge, central lit circle (with 50% illuminated crown) on dark background, and central dark circle (with 50% illuminated crown) on lit background (**Figure 1**).

### Tracking algorithm

The tracking algorithm involves three main steps: 1. background modelling; 2. object detection; and 3. object tracking (**Supplementary Figure 1**). First, the background was modelled using *N*_BG_ = 25 images acquired at uniform frequency during the whole experiment. Each of these images was converted to grey scale by selecting the red colour channel. Brightness was then re-scaled between 0 and 1, and a median blurring (window size *W*_blur_ = 9 *×* 9 for *E. gracilis*, and *W*_blur_ = 15 *×* 15 for *V. aureus*) applied to remove noise and smooth the contours of the objects present. Finally, these images were collapsed into a single output image using a pixel-wise median over the time samples. The resulting grey scale image were used to represent the background of the experiment (i.e., objects in the field of view that do not move and therefore should not be tracked).

After identifying the background, all images were then analysed in the order they were acquired. Each frame was first pre-processed using the same procedure as for the background modelling (i.e., channel selection, brightness scaling and blurring). Then, the foreground (i.e., the moving objects) was obtained using a pixel-wise subtraction between the current frame and the background. From the foreground, objects were detected using *brightness thresholding* (minimum brightness *b*_min_ = 85*/*255 for *E. gracilis*, and *b*_min_ = 90*/*255 for *V. aureus*). The identified object were then further selected according to their shape. Specifically, for each object we computed the area *a*, the perimeter *p* and the Polsby–Popper compactness measure [40], as *c* = (4*πa*)*/*(*p*^2^) ∈ [0; 1]. Only objects whose area and compactness fell within given ranges, [*a*_min_; *a*_max_] and [*c*_min_; *c*_max_] respectively, were retained (*a*_min_ = 175px, *a*_max_ = 1500px, *c*_min_ = 0.55, *c*_max_ = 0.9 for *E. gracilis*, and *a*_min_ = 800px, *a*_max_ = 6000px, *c*_min_ = 0.65, *c*_max_ = 1.0 for *V. aureus*). If the number of detected objects differed significantly (i.e., more than a factor δ = 25%) between consecutive frames, then the selection thresholds *a*_min_, *a*_max_, *c*_min_, *c*_max_ and *b*_min_ are iteratively relaxed or tightened by a factor *λ* = 2% until the number of objects is similar to the previous frame. If the condition on the number of detected objects is not satisfied after a maximum number of iterations, then the process is forcibly stopped.

The last step consisted of assigning IDs to the detected objects and ensuring these were consistent over time (i.e., an object ID is correctly propagated between frames). To do this, we considered the number *N*_*O*_ of all the previously detected objects and a generic one among them *j* ∈ *{*1,…, *N*_*O*_*}*. Its position at instant *k* was estimated as 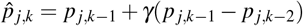,where *γ* = 0.9 is the inertia parameter. Moreover, its *inactivity I*_*j,k*_ was set to 0 if ID *j* was used at time *k* − 1, and incremented by one otherwise. Given the number *M*_*O*_ of objects detected in the current frame and their positions *p*_*i,k*_, their distance (in pixels) from object *j* is defined as 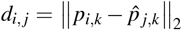.Similarly, their distance from the closest edge of the camera frame is denoted as *d*_edge,*i*_. Then, the cost of assigning ID *j* to the detected object *i* was defined as:

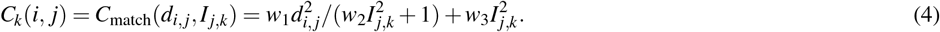

Moreover, the cost of assigning a new ID *j*^*′*^ ∈ *{N*_*O*_ + 1, …, *N*_*O*_ + *M*_*O*_*}* to *i* was defined as

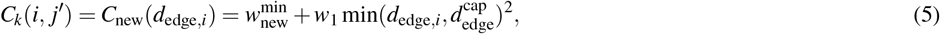

with the parameters set as follows, 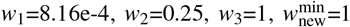 and 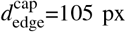 for *E*.*g*., and 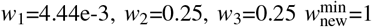 and 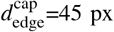 for *V*.*a*.. The Jonker-Volgenant optimization algorithm [41] was then applied on the resulting costs matrix 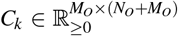 to solve the assignment problem, while minimizing the total cost. Specifically, each detected object was given an ID, either an existing or new one.

### Population kinetic data derivation

For each microorganism identified by the tracking algorithm, a multi step processing algorithm was further used to calculate the longitudinal velocity *v*_*k*_, and the angular speed *ω*_*k*_. First, each recorded trajectory was resampled at 0.5 s using linear interpolation, which compensated for possible jitter in the sampling time and provides missing data points. The interpolated trajectories were then smoothed using a moving average filter with window size *W*_ma_=3. After this refinement step, the trajectories having a duration shorter than the threshold *l*_min_=5 s were excluded to compensate for possible tracking errors. The remaining trajectories were then processed by extracting velocity vectors **v**_*k*_ ∈ *ℝ*^2^. These were computed by differentiating the trajectories using a backward Euler method. Specifically, we computed the velocity in px/s units and then converted it to µm/s units by multiplying by the pixel size *p* = 1.25 *µ*m for *E*.*g*. and *p* = 4.44 *µ*m for *V*.*a*.. From the velocity vector we then calculated the longitudinal velocity *v*_*k*_ = ∥**v**_*k*_∥_2_, and the angular velocity *ω*_*k*_ using,

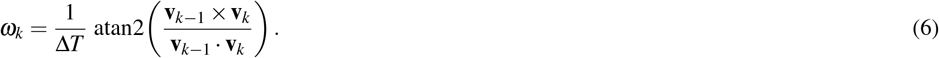

Finally, both *v*_*k*_ and *ω*_*k*_ were further smoothed using a moving average filter with window size *W*_ma_. Once the trajectory of every microorganism were processed, the average speed of each agent was computed and outliers of this distribution detected (see section below) using the threshold *m* = 1 for *E*.*g*. and *m* = 0.25 for *V*.*a*.. Among these outliers, those slower than the median (e.g., debris and dead microorganisms) were removed. A second outlier detection was performed on the instantaneous speed of the agents with threshold *m* = 1 for *E*.*g*. and *m* = 2 for *V*.*a*., and agents that at any time instant moved particularly fast (i.e., matching errors in the tracking algorithm) were removed. Finally, for each experiment, data obtained from the biological replicates were aggregated in a single dataset.

### Spatial data processing

Data from experiments with spatial inputs were processed to quantify the accumulation of microorganisms with respect to the light intensity across the sample. To performn this analysis, images were processed similarly as for the object detection. Specifically, they underwent channel selection, brightness scaling and median blurring. Finally, brightness thresholding was used to obtain a binary pixel mask. Unlike for object tracking, in this case we are not identifying individual objects, therefore, shape thresholding was not used. This allowed the analysis of images with a particularly high density of microorganisms, where detection of individual objects was not possible. The generated binary mask was then applied to the input light pattern so as to select only pixels corresponding to detected objects. The set of input values was binned (5 bins for the BCL experiment with *E. gracilis*, 3 bins for all other experiments), obtaining a counting of the selected pixels as a function of the input intensity. Similarly, pixel values of the whole input pattern were binned and counted. For each experiment this procedure was repeated for each biolgical replicate (3 for the BCL experiment with *E. gracilis*, 8 for other *E. gracilis* experiments, and 6 for experiments with *V. aureus*) and pixel counts from the replicates summed together. Finally, the resulting count of selected pixels was divided by the count of all the pixels and the result normalised to sum to one. This provided an approximation of the density of microorganisms as a function of the input light intensity. Similarly, for the simulated data the positions of the agents were used to compute their distribution as function of the input intensity.

### Error metrics

Several error metrics were used to assess the accuracy of our digital twins. Given a vector of data **x**, and a vector of estimations 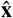,the quality of the estimation can be measured by computing the weighted Mean Absolute Percentage Error (wMAPE) [42], defined as

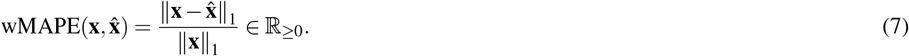

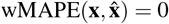 when the estimation perfectly fits the data. Conversely, 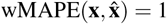 means that the estimates 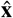 fit the data as good as the null vector. When validating the *E. gracilis* digital twin and assessing the generality of the phenotyping platform and model, we quantified the ability of the digital twins to reproduce the time evolution of experimental data by averaging the wMAPE computed for *v* and |*ω*|. More precisely, we defined

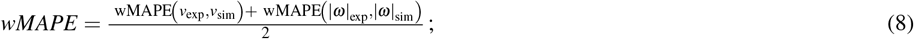

where |*ω*|_exp_(*v*_exp_) and |*ω*|_sim_(*v*_sim_) represent the median over the population of the absolute angular velocity (longitudinal velocity) in experiments and simulations, respectively.

To assess the ability of the digital agents to reproduce photoccumulation or photodispersion we quantified the difference between the experimental and simulated density distributions as functions of the input intensity *u*, denoted here as *ρ*_exp_(*u*) and *ρ*_sim_(*u*). We used the Total Variation Distance (TVD), defined for discrete distributions as,

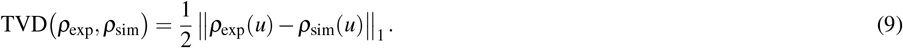

This metric is bounded between 0 and 1 and quantifies the difference between the distributions. Specifically, TVD (*ρ*_exp_, *ρ*_sim_) = 0 when the two distribution coincide, while TVD (*ρ*_exp_, *ρ*_sim_) = 1 when the two distributions have disjoint supports.

### Outliers detection

Given a data set *d* (vector or multidimensional array), and a positive threshold *m*, a data point *i* was classified as outlier if *d*_*i*_ *< Q*_1_ − *m* · IQR ∨ *d*_*i*_ *> Q*_3_ + *m* · IQR, where *Q*_*j*_ is the *j*-th quartile of *d* and IQR = *Q*_3_ − *Q*_1_ is the inter quartile range (IQR).

### Model parametrisation

Parametric identification was performed independently for *v* (Equation 1a) and *ω* (Equation 1b). This was possible as these processes are independent. For each, a multi-step *ad hoc* procedure was designed. First, we considered a specimen *i* of *E. gracilis*, and recalled that the evolution of its velocity *v*_*i*_ can be described according to Equation 2a as,

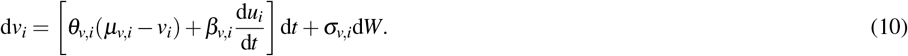

The identification procedure then begins by estimating the parameters that best fit the evolution of the average velocity of the population 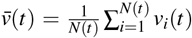.With this aim, we momentarily assumed that *θ*_*v,i*_ = *θ*_*v*_, *µ*_*v,i*_ = *µ*_*v*_ and *β*_*v,i*_ = *β*_*v*_, ∀*i*. Under these assumptions, and noticing that the input is spatially uniform (i.e., *u*_*i*_ = *u* ∀*i*), and acquiring *n* data points (i.e. 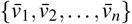 and *{u*_1_, *u*_2_, …, *u*_*n*_*}*), it is possible to use the Ordinary Least Squares method to find the values of *θ*_*v*_, *µ*_*v*_ and *β*_*v*_ that best fit the experimental evolution of the average velocity [43]. This set of parameters, approximating the evolution of the average velocity, is used as an initial guess and refined using the MATLAB GrayBox identification toolbox. This optimization is constrained with the assumption of stable dynamics (i.e., *θ*_*v*_ *>* 0), and imposing *β*_*v*_ *<* 0.

The refined parameters set fitting the evolution of mean speed of the population was then used as an initial guess for the MATLAB GrayBox identification toolbox to identify, for each specimen in the population, the values of *θ*_*v,i*_, *µ*_*v,i*_, *β*_*v,i*_ best fitting the time series of the speed of individual microorganism. Finally, we estimated the volatility as 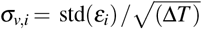,where *ε*_*i,k*_ = (*v*_*i,k*+1_ −*v*_*i,k*_)−[*θ*_*v,i*_(*µ*_*v,i*_ −*v*_*i,k*_) + *β*_*v,i*_(*u*_*k*_ −*u*_*k*−1_)] Δ*T*.

This same procedure was used to estimate the parameters describing the evolution of the longitudinal and angular velocity of *V. aureus*, described by the model in Equation 3.

To identify the parameters governing the angular velocity of *E. gracilis*, we recast Equation 2b so as to describe |*ω*|. Formally,

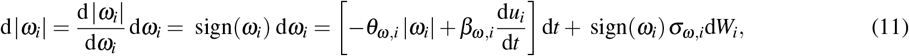

whose expected value then evolves as,

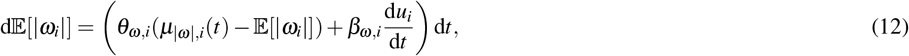

where *µ*_|*ω*|,*i*_(*t*) = *σ*_*ω,i*_ 𝔼 [sign(*ω*_*i*_) d*W*_*i*_]. Assuming that *µ*_|*ω*|,*i*_(*t*) = *µ*_|*ω*|,*i*_, we estimated 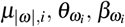 using the same procedure adopted for the longitudinal velocity. Finally, to estimate the value of *σ*_*ω,i*_ we computed the expected value of the steady state distribution of |*ω*_*i*_| in absence of an input. Specifically, *ω*_*i*_ will converge at steady state to 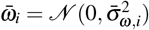,where 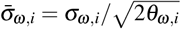.Hence, 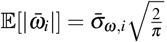.By noticing that 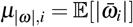, it is then possible to compute 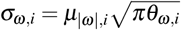.

Once all the parameters of all specimens in the populations were identified, we discarded all the individuals for which any of the parameters were classified as outlier with respect to rest of the population. This yielded a final dataset of *N*^*′*^ digital twins. The resulting distributions of parameters, for both *E. gracilis* (*N*^*′*^ = 182) and *V. aureus* (*N*^*′*^ = 19), are shown in **Figure 2h** and **Figure 6h**, respectively.

### Software modules

The software responsible for the data acquisition module was developed in Python and consists of two components. The first includes scripts running on Raspberry Pi OS that control the DOME, allowing for experiments to be automated, with little user input. The second component, developed in Python and executable on any laptop (Windows, macOS or Linux), reads the data acquired during the experiments, automatically tracks the microorganisms, and runs the algorithm to retrieve *v* and *ω*. The algorithm for the parametrization of the mathematical model and its validation is developed in MATLAB and integrated within SwarmSim. Unless otherwise stated, each simulation used a random selection of *N* digital twins extracted among those generated by our routine (*N* = 2000 for *E*.*g*. and *N* = 500 for *V*.*a*.). The selection probability was uniform across all digital twins and repetitions were allowed so that *N* could be larger than the number of digital twins *N*^*′*^ inferred. The trajectory of each digital twin was generated by integrating Equation 1 in time using an Euler–Maruyama algorithm (d*t* = 0.01 s). The simulations lasted for *T*_max_ = 180 s and the trajectories were sampled every Δ*T* = 0.5 s. Speed and angular velocity were computed from the simulated trajectories as above, but without any smoothing nor outlier detection and removal.

## DATA AVAILABILITY

**Supplementary Movie 1** shows a time-lapse of a *E. gracilis* population being exposed to a ‘BCL’ light pattern (letters were dark on a illuminated background). **Supplementary Movie 2** shows a comparison of the aggregated experimental data and digital twins of *E. gracilis* being exposed to a ‘BCL’ light pattern (letters were dark on a illuminated background). All experimental data from this study are available at: https://doi.org/10.5281/zenodo.13683455

## ACKNOWLEDGEMENTS

T.E.G. was supported by a Royal Society University Research Fellowship grant URF/R/221008, a Turing Fellowship from The Alan Turing Institute under EPSRC grant EP/N510129/1, BrisEngBio, a UKRI-funded Engineering Biology Research Centre under BBSRC grant BB/W013959/1, and the UKRI-funded Engineering Biology Mission Award CYBER under BBSRC grant BB/Y007638/1. The funders had no role in study design, data collection and analysis, decision to publish or preparation of the manuscript.

## AUTHOR CONTRIBUTIONS

T.E.G., M.d.B and A.G. conceived the project. T.E.G. and M.d.B. supervised the work. T.E.G. and A.G. designed the experiments. A.G. carried out all the experiments and developed the software components. M.d.B., A.G. and D.S. developed the model. A.G. and D.S. performed the data analysis and wrote the initial manuscript. All authors contributed to the interpretation of the results and editing of the manuscript.

