## Supplementary material for "Data-driven inference of digital twins for high-throughput phenotyping of motile and light-responsive microorganisms": Supplemena

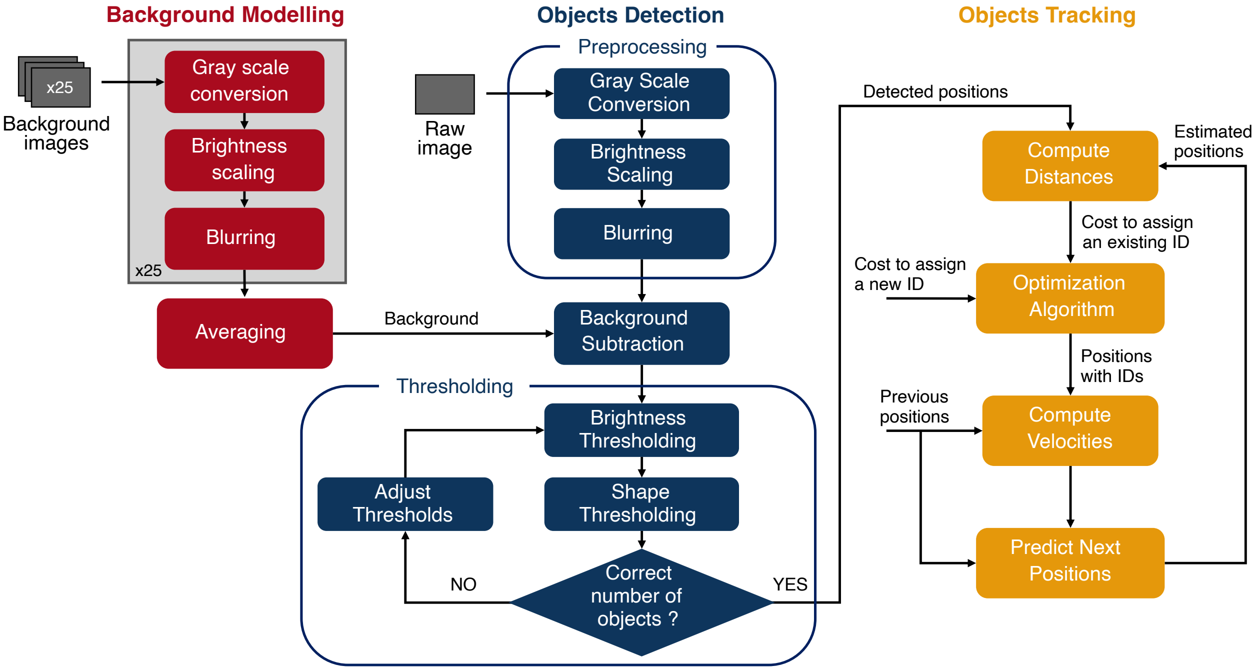

**Supplementary Figure 1: Overview of the tracking algorithm.** There are three core elements to the processing of raw microscopy images. First, background modeling is performed by generating a background mask (red section). Second, object detection is performed using a threshold-based approach (blue section). Finally, object tracking is carried out by making use of information from the previous frame (yellow section). See **Methods** for details about each step.

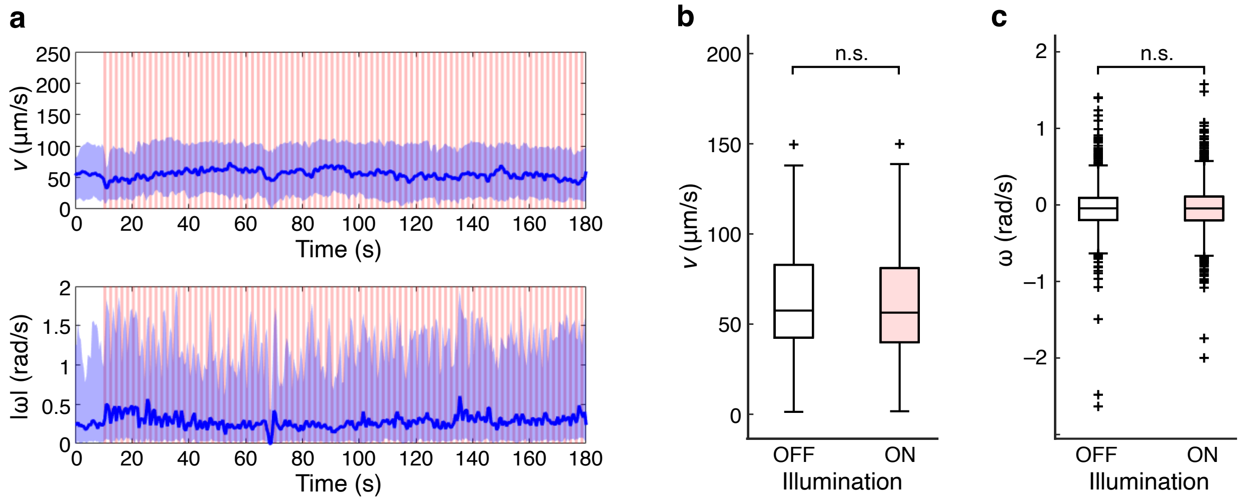

**Supplementary Figure 2: *E. gracilis* response to square waves with 1 s semi-period.** (**a**) *In vivo* time evolution of longitudinal velocity, $v$ (top), and absolute angular velocity, $|\omega|$ (bottom). Solid blue lines represent the median over the population and the shaded blue areas highlight the 10th–90th percentile range. Red shaded areas show the intervals when light inputs were present. (**b**) Comparison between the distributions of the average longitudinal velocity of the organisms in time when the light was off (white) and on (red). The distributions were compared using a Kolmogorov-Smirnov test (n.s.: not significant). (**c**) Comparison between the distributions of the average angular velocity of the organisms in time when the light was off (white) and on (red). The distributions were compared using a Kolmogorov-Smirnov test (n.s.: not significant).

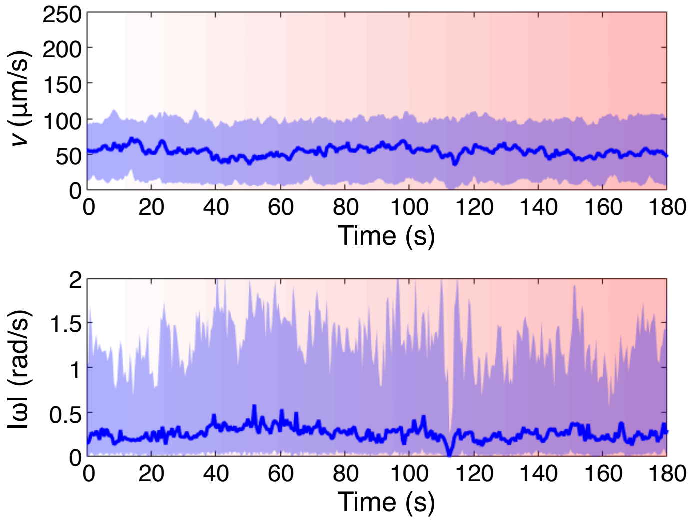

**Supplementary Figure 3: *E. gracilis* response to linearly increasing light intensity over time.** The time evolution of longitudinal velocity, $v$ (top), and absolute angular velocity, $|\omega|$ (bottom). Solid lines represent the median over the population while the shaded areas denote the $10^{th}$ to $90^{th}$ percentile range. The varying shade of red shows the increasing intensity of the light input (0% intensity at $t \leq 10$ s, to 100% intensity for $t \geq 170$ s).

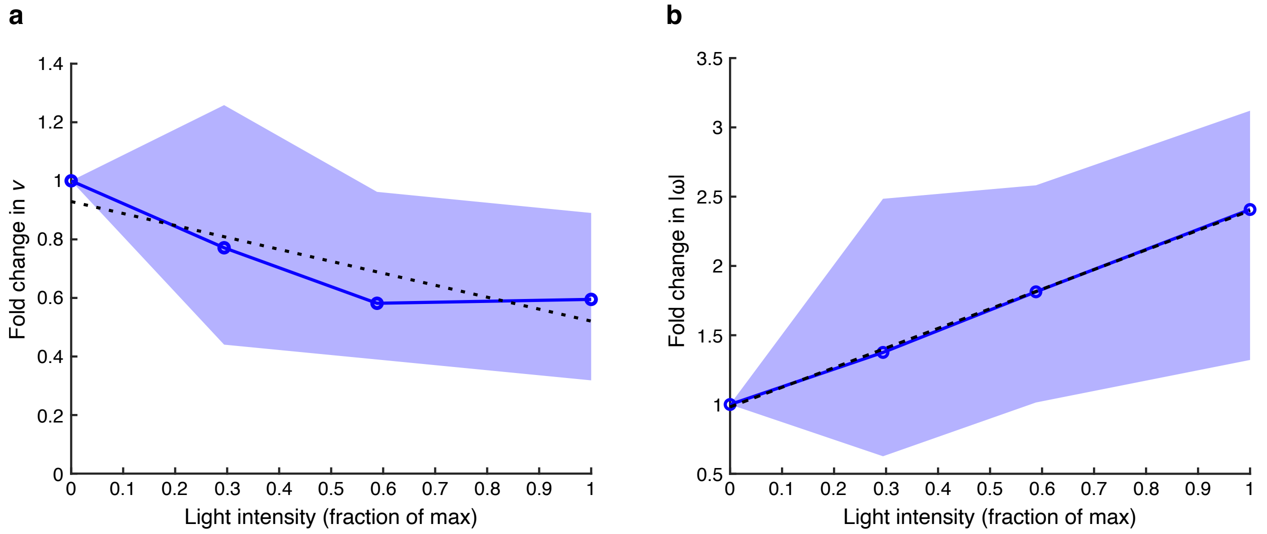

**Supplementary Figure 4: Response of *E. gracilis* to square waves of varying light intensity.** Fold change of longitudinal velocity, $v$, and absolute angular velocity, $|\omega|$, in dark and lit environment when a population of *E. gracilis* was stimulated with square waves having a semi-period of 60 s. (**a**) Fold-change of the mean longitudinal velocity as a function of the light intensity (shown as a fraction of maximum intensity). (**b**) Fold-change of the mean magnitude of the absolute angular velocity as a function of the light intensity (shown as a fraction of maximum intensity). In both panels, the fold-change is computed by normalizing the average $v$ or $|\omega|$ 12 s after the light switches on with respect to the the corresponding average observed 40 s before the light switches off. The blue solid line represent the average fold-change for 3 biological replicates, and the shaded blue area shows 25% to 75% of the distribution. The dashed black line represent a linear regression of the fold-change as a function of the light intensity.

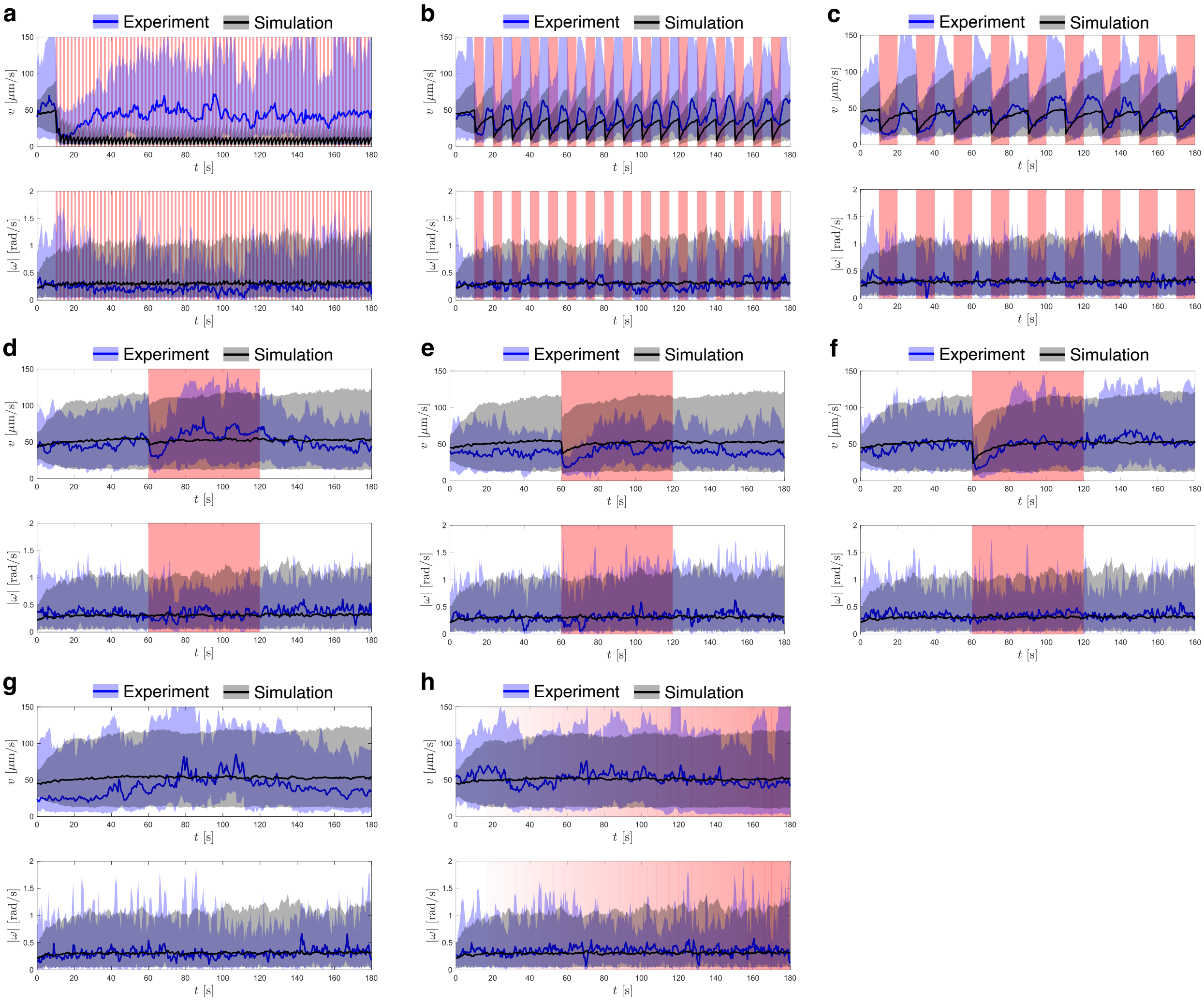

**Supplementary Figure 5: Validation of *V. aureus* model using time varying light inputs.** Comparisons of the time evolution of longitudinal velocity $v$, and absolute angular velocity $|\omega|$ (bottom), between experiments (combined biological replicates, blue) and model simulations (black) with different temporal light patterns. Solid lines represent the median over the population while the shaded areas highlight the 10th–90th percentile range. Red shaded areas highlight the intervals when light inputs were present. (**a**) Square wave light input, 1 s semi-period, 100% intensity. (**b**) Square wave light input, 5 s semi-period, 100% intensity. (**c**) Square wave light input, 10 s semi-period, 100% intensity. (**d**) Square wave light input, 60 s semi-period, 30% intensity. (**e**) Square wave light input, 60 s semi-period, 60% intensity. (**f**) Square wave light input, 60 s semi-period, 100% intensity (used for fitting of the model). (**g**) No illumination. (**h**) Linearly increasing light intensity input from 0% to 100%.

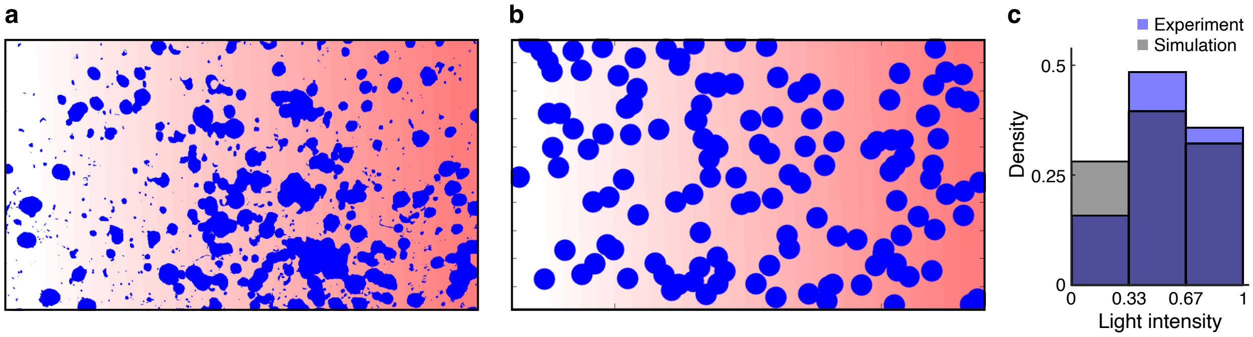

**Supplementary Figure 6: Photoaccumulation of *V. aureus* in a light gradient.** (**a**) Distribution of Volvox *in vivo* after a 180 s exposure to a static light gradient. (**b**) *In silico* distribution of Volvox after a 180 s exposure to a static light gradient from 0% (left) to 100% (right) intensity. (**c**) Density distributions of the microorganisms as a function of the normalized light intensity (simulations: dark grey; experiments: light purple).

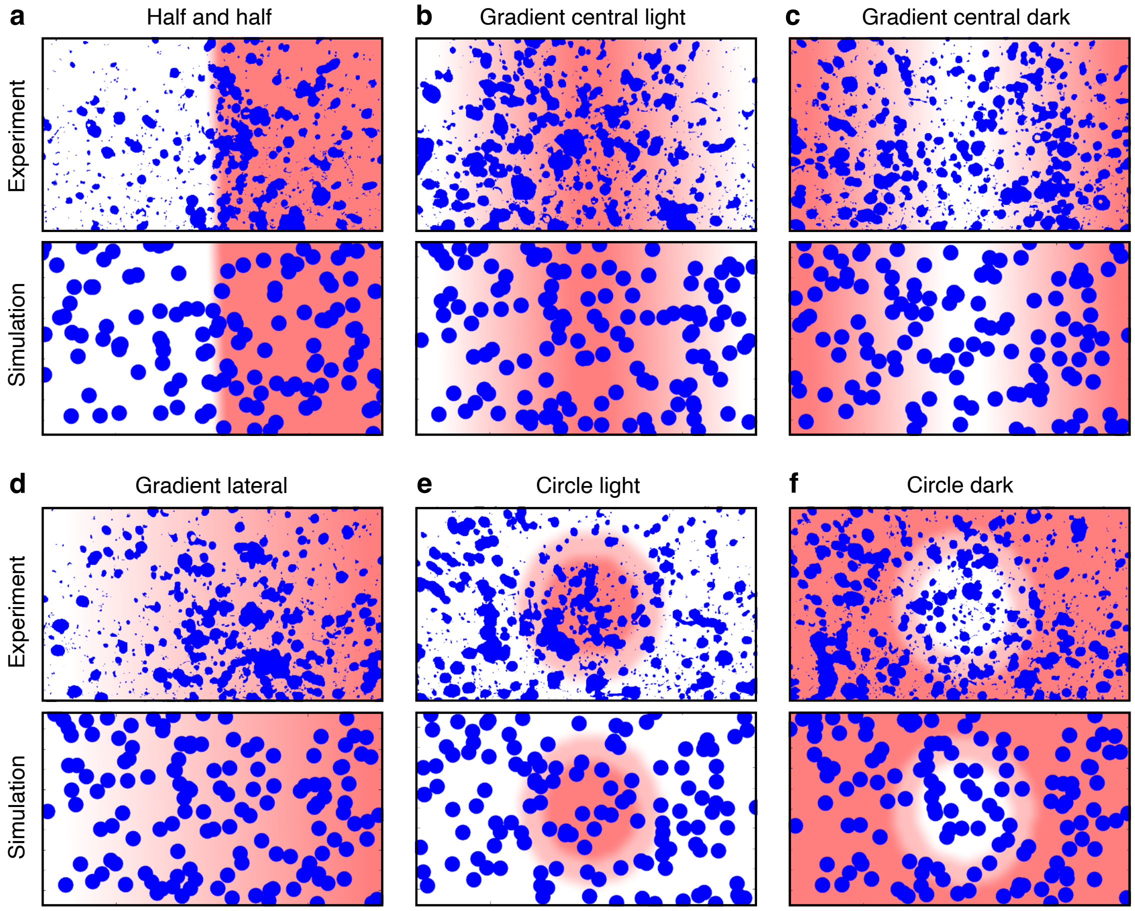

**Supplementary Figure 7: Validation of *V. aureus* model using spatially varying light inputs.** Snapshots from experiments and simulations after 180 s of exposure to a spatially patterned light input at 100% intensity. Red and white regions denote illuminated and dark/unilluminated areas, respectively. (**a**) Half and half light pattern. (**b**) Gradient from lit centre. (**c**) Gradient from unlit centre. (**d**) Lateral gradient dark to light (left to right). (**e**) Central lit circle and unlit background. (**f**) Central unlit circle and lit background.
